# Optimization and comparison of two methods for spike train estimation in an unfused tetanic contraction of low threshold motor units

**DOI:** 10.1101/2022.07.01.498432

**Authors:** Robin Rohlén, Christian Antfolk, Christer Grönlund

## Abstract

**Background:** Human movement is generated by activating motor units (MUs), i.e., the smallest structures that can be voluntarily controlled. Recent findings have shown imaging of voluntarily activated MUs using ultrafast ultrasound based on displacement velocity images and a decomposition algorithm. Given this, estimates of trains of twitches (unfused tetanic signals) evoked by the neural discharges (spikes) of spinal motor neurons are provided. Based on these signals, a band-pass filter method (BPM) has been used to estimate its spike train. In addition, an improved spike estimation method consisting of a continuous Haar wavelet transform method (HWM) has been suggested. However, the parameters of the two methods have not been optimized, and their performance has not been compared rigorously.

**Method:** HWM and BPM were optimized using simulations. Their performance was evaluated based on simulations and two experimental datasets with 21 unfused tetanic contractions considering their rate of agreement, spike offset, and spike offset variability with respect to the simulated or experimental spikes.

**Results:** A range of parameter sets that resulted in the highest possible agreement with simulated spikes was provided. Both methods highly agreed with simulated and experimental spikes, but HWM was a better spike estimation method than BPM because it had a higher agreement, less bias, and less variation (*p* < 0.001).

**Conclusions:** The optimized HWM will be an important contributor to further developing the identification and analysis of MUs using imaging, providing indirect access to the neural drive of the spinal cord to the muscle by the unfused tetanic signals.

## Introduction

The central nervous system initiates voluntary force production and human movement by delivering appropriate excitatory inputs to motor neurons in the spinal cord. Motor neurons are the last neural cells to process neural information, and they connect to muscles by a synaptic connection, the neuromuscular junction. A motor neuron and the muscle fibres it connects to form a so-called motor unit (MU). Each movement is generated by activating thousands of MUs, which can be seen as the smallest structures that can be voluntarily controlled.

Recent innovations have enabled imaging and analysis of MUs using magnetic resonance imaging (MRI) [1,2] and ultrafast ultrasound (UUS) [3–5]. These imaging techniques provide high-resolution spatiotemporal information regarding the kinematics of the muscle fibres that are part of a MU. Therefore, they provide complementary information to the gold standard techniques measuring MU activity in vivo, i.e., needle electromyography (EMG) invasively [6–8] and surface EMG non-invasively [9,10]. Both the MRI technique and the early UUS methods use electrical stimulation to generate muscle activity to image MUs. In contrast, a recently proposed UUS technique (MU-UUS) can identify single MUs in voluntary activations using a blind source separation algorithm of the velocity image sequences to extract spatiotemporal components [11] in which a subset corresponds to the MU [12–14]. Given this, MU-UUS provides estimates of MU territories in cross-section and the corresponding train of twitches evoked by the spinal motor neurons’ neural discharges (spikes). Suppose the discharge rates of the spinal motor neurons are low, resulting in an unfused tetanic contraction. In that case, it should be possible to identify the neural discharges (spike train) where a robust spike estimation method would lead to indirect access to the neural drive of the spinal cord via the spinal motor neurons to the muscles.

Previous studies involving the MU-UUS analysis have provided estimates of unfused tetanic signals of single MUs [11,12,14,15]. These unfused tetanic estimates have further been used to estimate their spike trains (neural discharges). The challenge for this spike estimation is that the unfused tetani of voluntary contractions consist of variable successive twitches [16–18]. For the MU-UUS analysis, a narrow band-pass filter has been used before applying peak detection to identify the time instants of the local maxima [11,12,15]. Although an optimized parameter set may have improved the performance, these studies did not focus on optimizing the parameters [11]. This spike estimation has been suggested to have lower performance for unfused tetanic signals with lower SNR [19], and the estimated spikes have been shown to vary with respect to the reference (experimental) spike [12]. For these reasons, an improved spike estimation method was suggested that consisted of the continuous Haar wavelet transform. It was motivated by its property of detecting abrupt transitions and oscillations in the signal as the twitch onset provides relatively large wavelet coefficients [19]. However, none of the parameters of the two methods (the band-pass filter method BPM and the Haar-wavelet method HWM) has been optimized, and the performance of the methods has not been compared rigorously.

This study aimed to optimize the parameters of BPM and HWM using simulations of unfused tetanic signals and to evaluate the optimized methods’ performance of the two methods in terms of 1) the rate of agreement, 2) the spike offset, and 3) the spike offset variability with respect to the simulated/reference spikes. The hypothesis was that the HWM should have higher performance for all three comparisons than BPM because the twitch onset is believed to be a more robust feature that varies less and foregoes the time instants of local maxima. The experimental performance of the optimized methods was evaluated using two datasets: one from human biceps brachii extracted using MU-UUS analysis and the second dataset from functionally isolated MUs in rat gastrocnemius.

## Methods

### Spike estimation

#### The band-pass filter method (BPM)

The BPM finds local maximums of the signal based on a band-pass filtered and standardised signal (Fig. 1). BPM consists of three steps:

**Figure 1.**
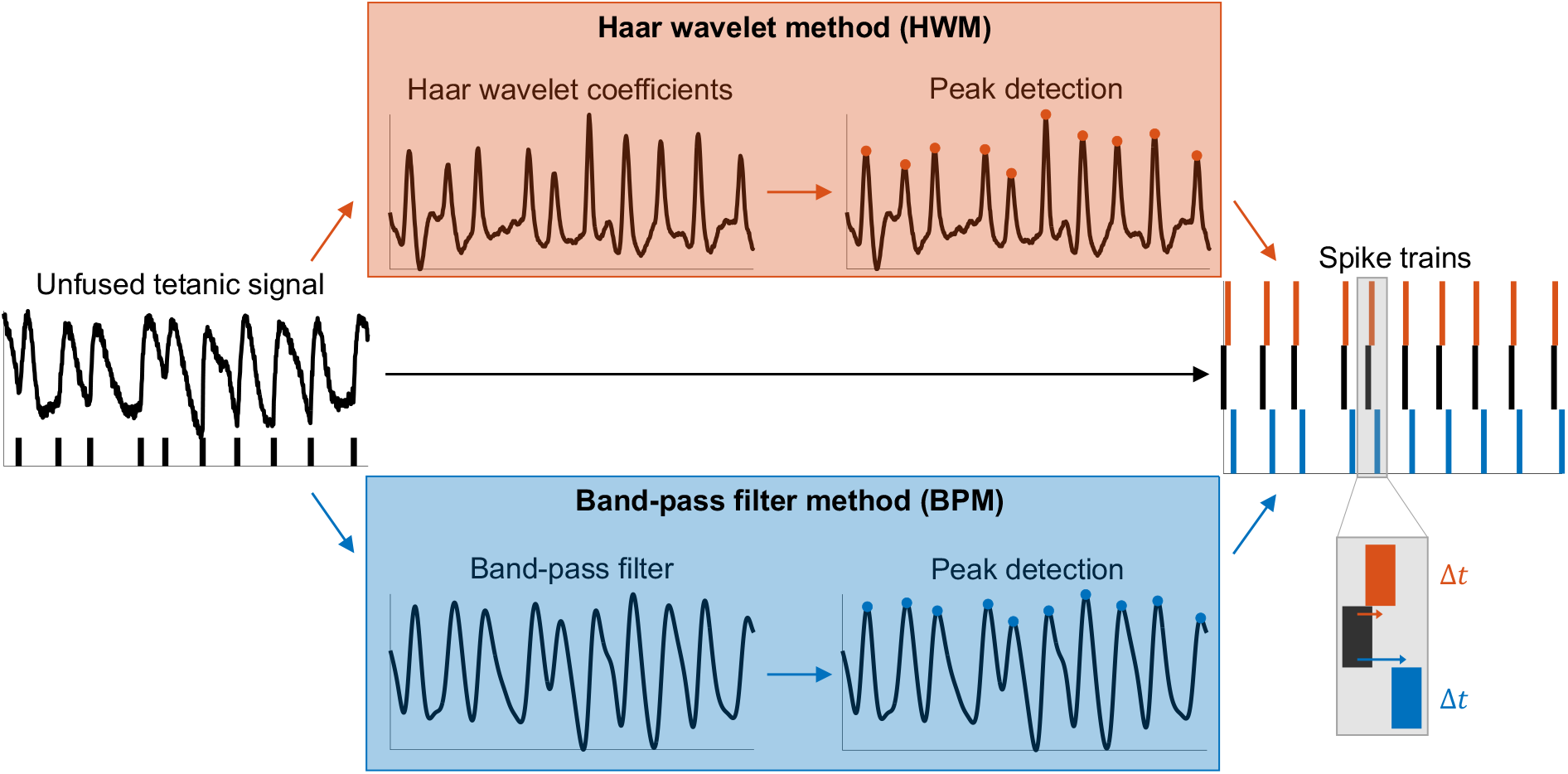
An illustration of the two methods used for estimating spike trains based on an unfused tetanic signal: the Haar wavelet method (HWM) and the band-pass filter method (BPM). The two methods consist of two steps: 1) filtering and 2) peak detection. Note that the HWM detects the steep gradient of each twitch, whereas the BPM detects the local amplitude maxima.

1. Applying a zero-phase Butterworth band-pass filter
2. Standardising the signal based on z-score
3. Finding the time instants of the local maxima of the signal using peak detection Step 1 requires three parameters: the filter order and the Butterworth filter’s low- and high-pass cut-off frequency. Step 3 requires two parameters: the minimal peak height of local maxima and the minimal peak distance between successive local maxima. After initial tests, we optimised three parameters: the low-pass cut-off, minimal peak height, and minimal peak distance (Table 1). That means the filter order and high-pass cut-off frequency were fixed and equal to 4 and 5 Hz, respectively. Note that the filter order was set to 2, but since we used a zero-phase digital filter that processes the signal in a forward and reverses fashion, the filter order is doubled.

**Table 1.** The parameters used in the optimization for the BPM and the HWM.

|  | **BPM** | **HWM** |
| --- | --- | --- |
| **Filtering** |  |  |
| Filter order | 4* | - |
| Low-pass cut-off (Hz) | (12, 14, ..., 38) | - |
| High-pass cut-off (Hz) | 5 | - |
| Scale** | - | (12, 14, ..., 38) |
| Pseudo-frequencies (Hz) |  | (26, 28, ..., 83) |
| **Peak detection** |  |  |
| Min peak height (z-score) | (0.02, 0.04, ..., 0.42) | (0.60, 0.62, ..., 1.00) |
| Min peak distance (ms) | (10, 11, ..., 40) | (10, 11, ..., 40) |

BPM = band-pass filter method, HWM = Haar-wavelet method. The filtering is based on a Butterworth filter and a zero-phase forward and reverse digital infinite impulse response filtering. \* The filter order is double (order 2x2=4) because of the zero-phase filter in forward and reverse directions. \*\* The simulated data had a sample rate of 1 kHz and based on these values the pseudo-frequencies for the Haar wavelet were approximated.

#### The Haar-wavelet method (HWM)

In contrast to BPM, HWM finds the signal’s rises (steep gradients) based on a continuous Haar wavelet transform (Fig. 1). The motivation for using the Haar wavelet is based on detecting abrupt transitions and oscillations in the signal providing large wavelet coefficients. In addition, smooth signal features produce large wavelet coefficients at pseudo-frequencies where the oscillation in the wavelet correlates best with the signal feature. Also, the Haar wavelet has compact support (non-zero only on a small sub-interval) at all pseudo-frequencies; its continuous wavelet transform coefficients are not affected by signal values at samples far away.

HWM consists of three steps:

1. Computing the continuous Haar wavelet transform of the signal at a specific pseudo-frequency (or scale)
2. Standardising the continuous wavelet transform coefficients based on z-score
3. Finding the time instants of the local maxima of the wavelet coefficients using peak detection

Step 1 requires one parameter: the pseudo-frequency, which can be approximated based on a scale *s* and the centre frequency *F*_*c*_ of the Haar wavelet by 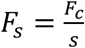. So, a scale is inversely proportional to the pseudo-frequency. Step 3 is the same as for the BPM. We optimized three parameters: pseudo-frequency, minimal peak height, and minimal peak distance (Table 1).

### Simulated data

We simulated unfused tetanic signals to optimize the parameters of the spike estimation methods (BPM and HWM) and evaluate their performance. This study’s simulated data was based on simulating 1) a spike train and 2) twitches that vary, which has been explained in other studies for voluntary contractions [16–18] for each spike in the spike train.

First, the time instants of the spikes within a given spike train were simulated using a Gaussian renewal process (Fig. 2A) [20]. Next, the mean was set to be equal to the mean inter-spike interval (ISI), i.e., the inverse of the mean firing rate and a standard deviation equal to the mean ISI times the coefficient of variation in ISI (ISI CV). Finally, we chose three firing rates (8, 12, and 16 Hz) [12,21] and different ISI CVs (5, 20, and 40%) [22] to simulate physiological firing rates at low force levels (low threshold MUs).

**Figure 2.**
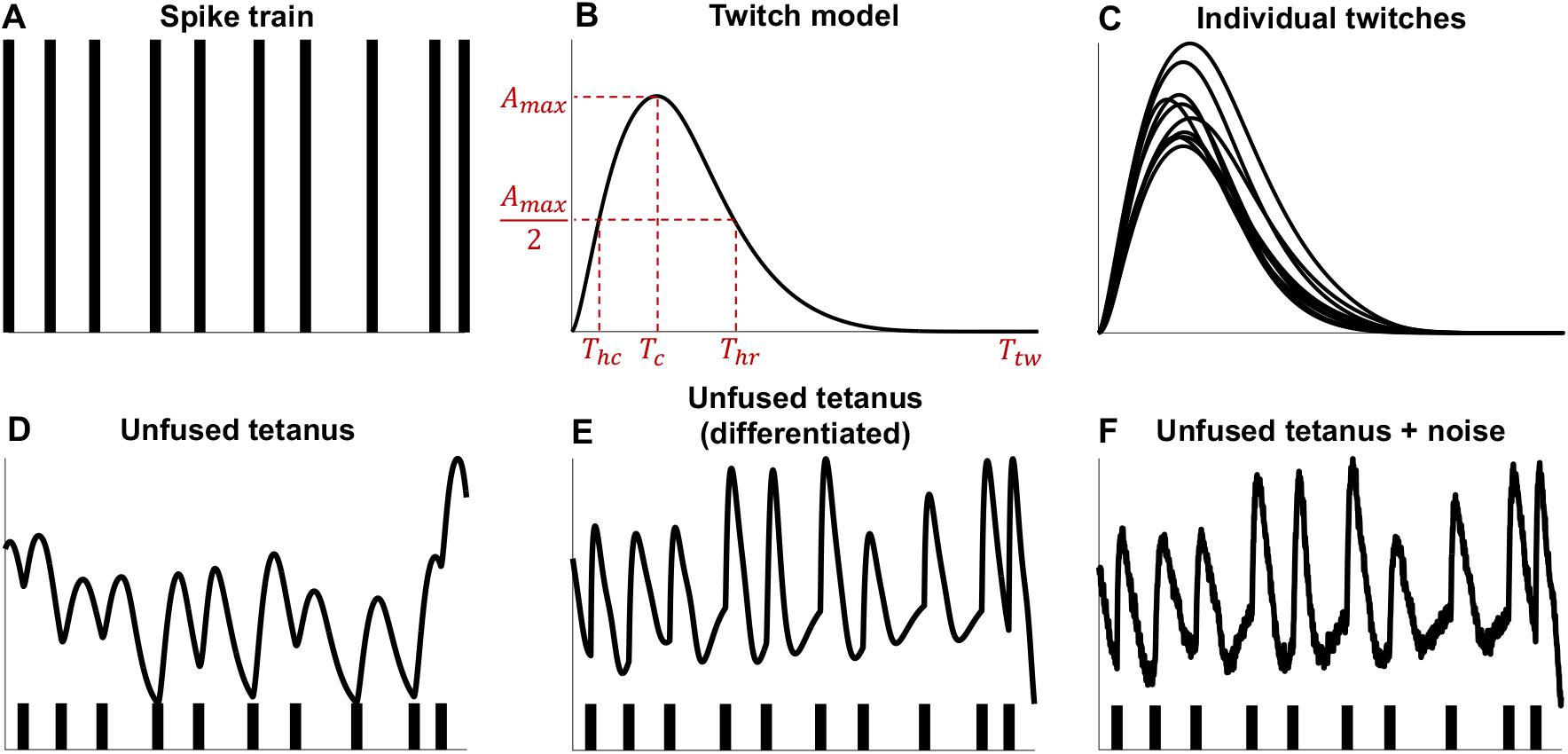
Simulating unfused tetanic signals using a spike train and a six-parameter twitch model. **A**. Time instants (spikes, black vertical lines) for each twitch were simulated using a Gaussian renewal process. **B**. A twitch (black line) is generated by its six parameters (red dashed lines) of the model are: T_hc_ – the half-contraction time (ms); T_c_ – the contraction time (ms); T_hr_ – the half-relaxation time (ms); T_tw_ – the twitch duration (ms); and A_max_ – the maximal amplitude (arbitrary unit, a.u.). The sixth parameter, which is known, is the time instant of spike either simulated or from reference signal (electromyography). **C**. The parameters of the successive twitches were randomly sampled from a uniform distribution resulting in varying shapes of the individual twitches with a bell-shaped form. **D**. The unfused tetanic signal was obtained by summating the twitches in **C** at their respective time instants in **A**. Note that the time instants for the first and last twitch have been omitted to make the first and last value of the signal to be non-zero. **E**. Unfused tetanus in **D** were differentiated with respect to time, providing unfused tetanus concerning the rate of change. For example, suppose tetanus in **D** is in displacement. In that case, tetanus in **E** is in velocity, where each velocity twitch has a positive contraction and a negative relaxation. **F**. Then, Gaussian noise was added to achieve a certain signal-to-noise (SNR) level. Here, the mean ISI was 125 ms (8 Hz firing rate) with a coefficient of variation of 20% and SNR equal to 20 dB.

Second, a six-parameter twitch model [18] was used for simulating the twitches. These parameters are the lead time (*T*_*lead*_), the half-contraction time (*T*_*hc*_), the contraction time (*T*_*c*_), the half-relaxation time (*T*_*hr*_), the twitch duration (*T*_*tw*_), and the maximal amplitude (*A*_*max*_). The sixth parameter, considered known, is the time instant of the spike that precedes the lead time and initiates the twitch (see the previous paragraph). The analytical equations are presented in Raikova et al. (2008) [18], and the five parameters are illustrated in Fig. 2B. The twitch parameters were sampled from a uniform distribution with pre-defined ranges of the parameters suitable for low threshold MUs at low force levels [19] (Table 2). The lead time (*T*_*lead*_) can be interpreted as the time from spike to twitch onset. The stimuli can be from electrical stimulation or a neural discharge, as in voluntary contractions. So, the lead time is related to the electromechanical delay and is assumed to be constant within a contraction [23]. In this paper, we set the lead time to be zero since a different value only introduces shifts in the locations of the estimated spikes, rate of agreement threshold (see Parameter optimization), etc.

**Table 2.** The simulation parameters of the unfused tetanic contractions.

| Parameter | Value(s) |
| --- | --- |
| **Whole unfused tetanus** |  |
| Twitches/spikes (per simulation) | 100 |
| Repeated simulations | 100 |
| SNR (dB) | (10,20,30) |
| **Spike train parameters** |  |
| Firing rate (Hz) | (8,12,16) |
| ISI CV (%) | (5,20,40) |
| **Twitch parameters** |  |
| $T_{lead}$ (ms) | Uniform(0,0) |
| $T_{hc}$ (ms) | Uniform(20,25) |
| $T_c$ (ms) | Uniform(50,75) |
| $T_{hr}$ (ms) | Uniform(100,130) |
| $T_{tw}$ (ms) | Uniform(300,350) |
| $A_{max}$ (n.u.) | Uniform(40,70) |

SNR = signal-to-noise ratio, ISI = inter-spike interval, CV = coefficient of variation. Uniform(a,b) = the uniform distribution with parameter a and b.

After simulating the twitches (Fig. 2C), two additional twitches were simulated and used for the summation of twitches (the first and last twitch). The next step is to sum each twitch to the respective spike resulting in unfused tetanus. After the summation, we removed the contributions from the first and last twitch to achieve a stable signal, mimicking the experimental situation of MU-UUS, where one records muscle contractions in the middle of a contraction (Fig. 2D). Then, the unfused tetanus was differentiated with respect to time (Fig. 2E). So, if the twitch is in terms of displacement, the differentiated twitch is in terms of velocity. If the twitch is in terms of force, the rate of force development is the yank [24]. There are two motivations for using the derivative of unfused tetanus with respect to time. First, the time derivative of unfused tetanus is often stabilized around its mean value because the positive and negative parts of a contraction approximately cancel each other. In contrast, unfused tetanic contractions (in terms of force- or displacement twitches) are non-stabilized signals, where the signal’s shape depends on the firing information and MU properties. Second, the method for identifying single MUs using ultrafast ultrasound (MU-UUS) decomposes velocity image sequences into spatiotemporal components where the temporal part of the component is in terms of velocity.

The final step in the simulations was to add Gaussian noise to the unfused tetanic signal, so the signal-to-noise ratio (SNR) was 10, 20, or 30 dB (Fig. 2F). In this way, each simulation was generated by combining the three SNR values (30, 20, and 10 dB), ISI CV values (5, 20, 40%), and the firing rates (8, 12, 16 Hz) resulting in 27 (3×3×3) combinations. Each combination generated a total of 100 twitches within an unfused tetanic signal. We repeated this procedure 100 times for each combination, which resulted in 27×100=2,700 unfused tetanic signals and 2,700×100=270,000 individual twitches. A summary of the simulation parameters used can be found in Table 2.

### Experimental data

#### Unfused tetanic signals from ultrasound imaging on human biceps brachii in vivo

Retrospectively included from a previous study, three estimated unfused tetanic signals from three healthy subjects (28.3 ± 0.6 years old, two women and one man) were used to test the optimized spike estimation methods (BPM and HWM). These signals were two seconds long and a subset of those obtained in a previous study from the cross-section of the biceps brachii at low force levels (about 1% of maximum voluntary contraction) using the MU-UUS analysis [12]. The subjects performed low force isometric elbow flexion (90 degrees) and supination of their lower arm as a physician inserted a concentric needle into the biceps brachii muscle and iterated until the physician obtained clear EMG signals. Then the ultrasound probe was positioned proximal to needle insertion such that the image plane captured the needle tip. A concentric needle electrode was used simultaneously as the ultrafast ultrasound to record single MU action potentials for each unfused tetanic signal. The ultrasound and EMG systems were synchronized with the sampling rates of 2 kHz (radio frequency data of the ultrasound system) and 64 kHz (needle EMG). The needle EMG data were used to identify the spike train of the MUs as a reference to the estimated spike trains from BPM and HWM. The subjects gave informed consent before the experimental procedure. The project conformed to the Declaration of Helsinki and was approved by the Swedish Ethical Review Authority (2019-01843). See Rohlén et al. (2020) [12] for detailed information about the experimental procedure.

#### Unfused tetanic signals from electro-stimulation on functionally isolated motor units in rat gastrocnemius

Retrospectively included from a previous study, eighteen unfused tetanic signals were measured from five adult female Wistar rats under pentobarbital anaesthesia. The medial gastrocnemius muscle and the respective branch of the sciatic nerve were partly isolated from surrounding tissues, while other muscles of the hindlimb innervated by the sciatic nerve were denervated. Laminectomy over segments L2-S1 was performed, and the L5 and L4 ventral roots were cut proximally to the spinal cord and split into very thin bundles of axons. The animals were immobilized in a steel frame, and the operated hind limb and the spinal cord were covered with paraffin oil. The Achilles tendon was connected to an inductive force transducer to measure the contractile force under isometric conditions and stretched to the passive force of 100 mN [25]. The thin bundles of axons were electrically stimulated with suprathreshold rectangular pulses (amplitude up to 500 mV, duration 100 µs). The all-or-none character of the evoked twitch contractions and action potentials were used as criteria for a single MU isolation [26,27].

The MU action potentials were recorded with two non-insulated silver wire electrodes (150 μm in diameter) inserted through the middle part of the muscle, perpendicular to its long axis. The MU action potentials were amplified by a multi-channel preamplifier (World Precision Instruments, model ISO-DAM8-A) with a ground electrode inserted into the muscles of the opposite hind limb. The force and EMG signals were stored using a 12-bit analogue-to-digital converter (RTI-800) with a sampling rate of 1 kHz and 10 kHz, respectively.

To ensure that the included unfused tetanic signals were from slow MUs, the MUs were classified based on the shape of 40 Hz unfused regular tetanus. The signals were classified as fast-twitch if a so-called “sag” was observed, whereas “non-sagging” signals were classified as slow-twitch characteristics [28,29]. For three rats, the stimulation frequencies were 10, 12.5, 14.3, and 16.6 Hz (ISI equal to 100, 80, 70, and 60 ms), while in two rats, the stimulation frequencies were 10, 12.5, and 14.3 Hz. The intervals between the individual spikes were randomly set at values in the mean ISI ± 50% range.

All experimental procedures followed the European Union animal care guidelines and the principles of Polish Law on The Protection of Animals and were approved by the Local Bioethics Committee. For details about the surgical procedure, including video, on the functional isolation of MUs, see Drzymała-Celichowska and Celichowski (2020) [30].

### Parameter optimization

The parameter optimization was based on the rate of agreement (RoA) between the estimated and simulated firings in simulated data, which has been used in other studies for the same or similar purposes [11,12,31]. The RoA metric is defined as 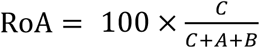, where *C* is the number of estimated spikes within an estimated spike train that was correctly identified, *A* is the number of false spikes, and *B* is the number of missed spikes. We set the tolerance interval for spike acceptance for the simulations (and the experimental data for voluntary human contractions) to be between 5 and 45 ms for BPM. For HWM, it was set to be between -5 and 25 ms. These thresholds were based on sensitivity analysis (Fig. S1, Supplementary Material), which infers the likely time instants of the amplitude maximum of each twitch (relevant for BPM) and positions of the rise time (relevant for HWM) for the simulated signals including added noise. For the unfused tetanic signals of the experimental rat data, we adjusted the tolerance interval for spike acceptance to be between 0-30 ms and 20-60 ms, respectively. This modification was because the data was in terms of force and not the yank (the derivative with respect to time), causing the local maxima of each twitch to be much larger and spread out. In contrast, the twitch’s rise only shifts slightly (see Fig. 5).

The optimization procedure worked as follows: for each simulated unfused tetanic signal and each spike estimation method, we calculated the RoA for all combinations of the parameters to be optimized (Table 1). The result (for each simulated signal and method) resulted in a 3D matrix of RoA values, with each dimension having the size of the parameter range. For example, the 3D matrix of RoA values for one simulated signal for HWM was 14×21×31. The first dimension stands for 14 different pseudo-frequencies and the second for 21 different minimal peak height values. The third dimension stands for 31 different minimal peak distance values. This procedure continued for all simulated signals (2,700 in total, each with 100 twitches and 100 spikes), which resulted in a 4D matrix of RoA values for each method where the fourth dimension equalled 2,700. Then we calculated the mean values across the 4^th^ dimension to get a 3D matrix with mean RoA values with each dimension corresponding to the range of parameter values associated with the values presented in Table 1. Finally, we identified the parameters providing the highest average RoA values.

### Performance evaluation

To evaluate the methods’ performance in terms of 1) the rate of agreement, 2) the spike offset, and 3) the spike offset variability with respect to the simulated/reference spikes, simulations and experimental data were considered. For the simulated data, we considered three cases of increasing complexity, and each case consisted of 50 unfused tetanic signals with 100 twitches each. The three simulation cases were to test the methods’ performance with respect to different simulation parameters: the SNR, ISI CV, and firing rate. The first case included a firing rate of 8 Hz, an ISI CV of 5%, and an SNR of 30 dB. The second case included a firing rate of 12 Hz, an ISI CV of 20%, and an SNR of 20 dB. The third case included a firing rate of 16 Hz, an ISI CV of 40%, and an SNR of 10 dB. The first quantification of performance was based on RoA. For spike offset (spike delta) and spike offset variability (spike delta variation), the estimated spikes that were correctly classified (*C*) in the RoA calculation were considered. The RoA values, spike deltas, and spike delta variations were used for statistical analysis between the two methods.

The experimental performance of the optimized methods was evaluated using two experimental datasets, with the first dataset being the human biceps brachii data and the second dataset being the rat gastrocnemius data. Before spike train estimation, the signals of the rat dataset were high-pass filtered with a 3 Hz zero-phase Butterworth filter with order 6 (2×3 due to forward-reverse). Similar to the simulation performance evaluation, RoA was calculated for each of the 21 signals and the correctly classified spike deltas, and spike delta variations were considered. Before using these features for statistical analysis, the averaged spike delta of dataset 1 was adjusted (for each of the three signals) such that the mean was located around 4 ms for HWM and around 20 ms for BPM. These values are selected based on the offset in the simulations. This adjustment makes the comparison of spike estimation variation possible because the electromechanical delay may be subject-dependent [32].

### Statistics

The statistical analysis of the difference between the methods’ performance (HWM and BPM) in RoA was performed using Wilcoxon signed-rank test to assess the median differences of paired RoA values. The same test was used to test the differences between the spike deltas. The difference in spike delta variation between the spike estimations of the two methods was tested using Levene’s test [33]. Note that the number of unfused tetanic signals in dataset 1 (voluntary human contractions) was so few (n=3) that testing the difference between the two methods was not feasible. Therefore, those results were presented but not subject to statistical testing. The data processing and analysis were performed using MATLAB (2022a, MathWorks, Natick, MA, USA).

## Results

### Simulations: parameter optimization

The five largest mean RoA values for HWM with the pseudo-frequencies between 45 and 71 Hz ranged between 97.3-97.7%, with a standard error of about 1.4% (Fig. 3A). For these pseudo-frequencies, the optimal minimal peak heights and minimal peak distances were between 0.88-0.96 and 15-25 ms, respectively (Fig. 3B-F). The relative positions of the RoA vs the pseudo-frequency were about the same across different simulated parameters except for the 8 Hz with 10 dB SNR, where the larger pseudo-frequencies had lower mean RoA (Fig. S2, Supplementary Material). Given these results, we selected the parameter set for HWM to be the following: the pseudo-frequency, minimal peak height, and minimal peak distance to be 45 Hz, 0.94 n.u., and 25 ms, respectively.

**Figure 3.**
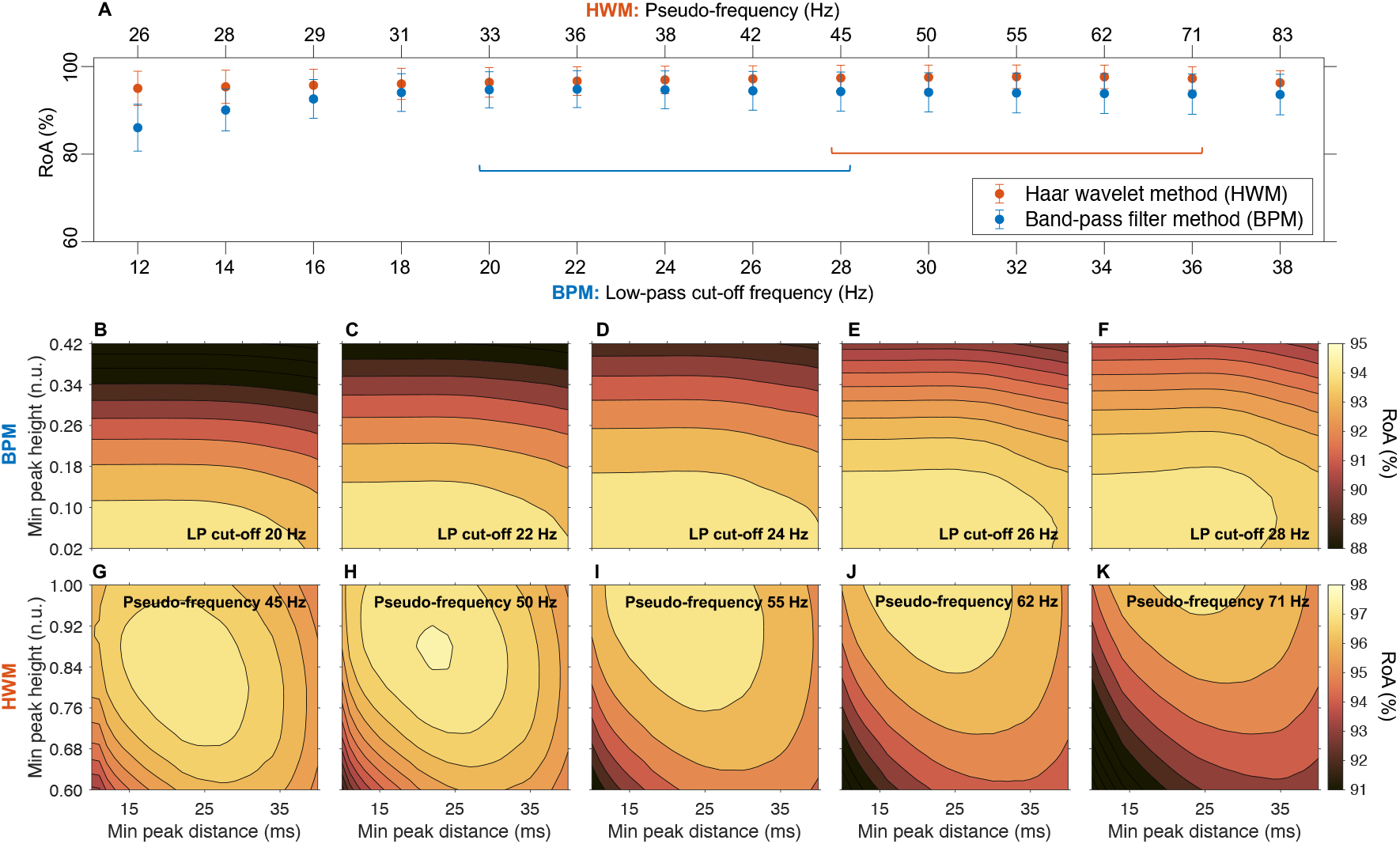
Parameter optimization. Quantifying the optimal parameters for each method using simulated data and the rate of agreement (RoA) metric. A total of 270,000 twitches based on different firing rates (8-16 Hz), inter-spike interval coefficient of variations (5-40%), and signal-to-noise ratios (10-30 dB) were considered. **A**. The mean RoA with respect to different pseudo-frequencies (orange dots) concerning the Haar-wavelet method (HWM) and different low-pass cut-off (blue dots) parameters concerning the band-pass filter method (BPM). The coloured horizontal lines show the five highest RoA values for each method’s parameter: 20 to 28 Hz for BPM (**B**-**F**) and 45 to 71 Hz for HWM (**G**-**K**). These findings suggest that the BPM parameters that maximize RoA are a low-pass cut-off ranging between 20-28 Hz, a minimal peak height ranging from 0.02-0.10, and a minimum peak distance ranging from 10-40 ms. The HWM parameters that will maximize RoA are a pseudo-frequency ranging from 45-71 Hz, a minimal peak height ranging from 0.88-0.96, and a minimum peak distance ranging from 15-25 ms.

The five largest mean RoA values for BPM with the low-pass cut-off between 20 and 28 Hz ranged between 94.3-94.8%, with a standard error of about 2.2% (Fig. 3A). For these low-pass cut-offs, the optimal minimal peak heights and minimal peak distances were between 0.02-0.10 and 10-40 ms, respectively (Fig. 3G-K). The relative positions of the RoA vs the low-pass cut-off were about the same across different simulated parameters (Fig. S2, Supplementary Material). Given these results, we selected the parameter set for BPM to be the following: the low-pass cut-off, minimal peak height, and minimal peak distance to be 24 Hz, 0.10 n.u., and 25 ms, respectively.

### Performance evaluation – Simulations

The RoA associated with HWM and BPM were high for all cases: case 1 (99.9 ± 0.2 and 97.0 ± 1.6), case 2 (98.4 ± 1.3 and 95.9 ± 2.0), and case 3 (90.6 ± 2.8 and 88.4 ± 2.7) (Table 3). There was a difference in RoA between the two methods across all cases (*p* < 0.001). Moreover, the average spike delta for HWM and BPM was around 4 ms and 20 ms for all three cases (Table 3, Fig. 4). There was a significant difference between the methods’ difference in spike delta (*p* < 0.001) and spike delta variation (*p* < 0.001) across all three cases.

**Table 3.** Performance evaluation based on simulated and experimental signals using the rate of agreement (RoA), the spike delta (Δt), and the spike delta variation between estimated and simulated/reference spikes.

|  | Simulations<br>Case 1 | Simulations<br>Case 2 | Simulations<br>Case 3 | Voluntary<br>contractions | Evoked<br>contractions |
| --- | --- | --- | --- | --- | --- |
| # signals | 50 | 50 | 50 | 3 | 18 |
| # spikes/twitches | 5,000 | 5,000 | 5,000 | 58 | 720 |
| Firing rate (Hz) | 8 | 12 | 16 | 11.4 | 14.4 |
| ISI CV (%) | 5 | 20 | 40 | 13.4 | 30.6 |
| **HWM** |  |  |  |  |  |
| RoA (%) | $99.9 \pm 0.2$ | $98.4 \pm 1.3$ | $90.6 \pm 2.8$ | $96.3 \pm 3.2$ | $96.5 \pm 5.6$ |
| $\Delta t$ (ms) | $4.1 \pm 2.6$ | $4.3 \pm 2.7$ | $4.6 \pm 3.5$ | $4.9 \pm 1.7$ | $12.7 \pm 1.8$ |
| **BPM** |  |  |  |  |  |
| RoA (%) | $97.0 \pm 1.6$ | $95.9 \pm 2.0$ | $88.4 \pm 2.7$ | $87.2 \pm 10.1$ | $88.0 \pm 7.9$ |
| $\Delta t$ (ms) | $20.3 \pm 2.8$ | $19.7 \pm 4.0$ | $19.1 \pm 6.6$ | $18.5 \pm 3.7$ | $32.5 \pm 7.0$ |

**Figure 4.**
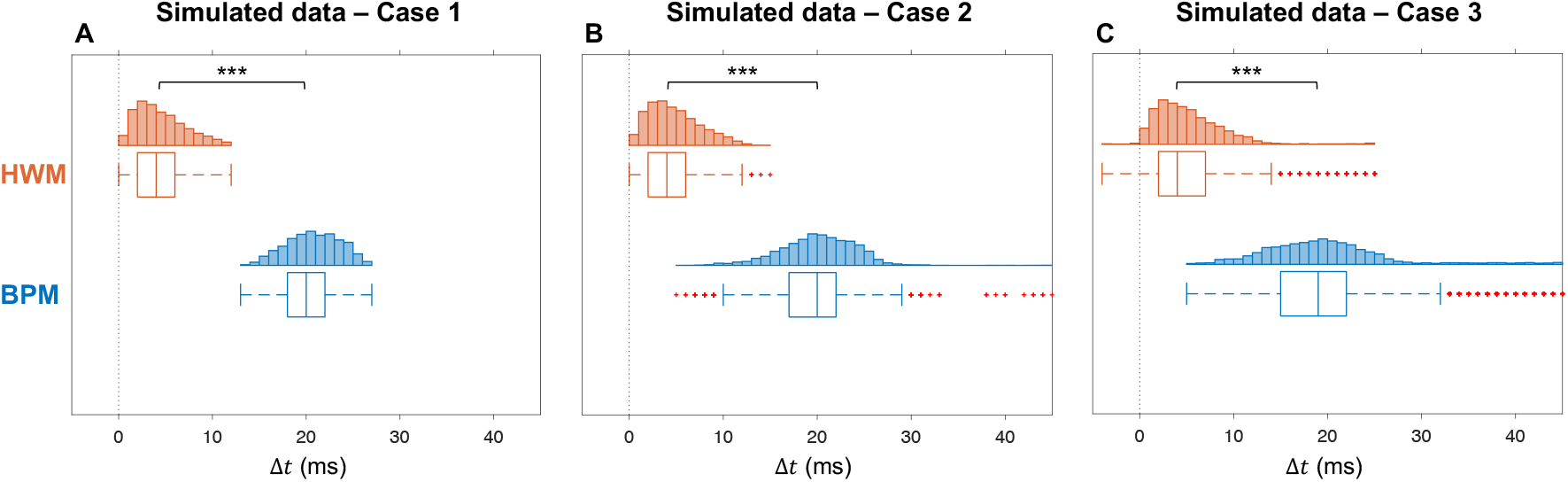
Performance evaluation – Simulations. Quantifying the methods’ delay (spike delta) and variation (spike delta variation) between the estimated and the simulated spikes in three cases of complexity where the simulated data consisted of 5,000 spikes and subsequent twitches. **A**. Case 1: The firing rate was equal to 8 Hz, the inter-spike interval coefficient of variation (ISI CV) equal to 5%, and the signal-to-noise ratio equal to 30 dB. **B**. Case 2: The firing rate was equal to 12 Hz, the inter-spike interval coefficient of variation (ISI CV) equal to 20%, and the signal-to-noise ratio equal to 20 dB. **C**. Case 3: The firing rate was equal to 16 Hz, the inter-spike interval coefficient of variation (ISI CV) equals 40%, and the signal-to-noise ratio equals 10 dB. For all cases, the spike train estimates of HWM were centred at 4 ms from the simulated spikes, while the BPM was centred at 20 ms. The estimated spikes of HWM had less spike delta and spike delta variation than the estimated spikes by BPM (p < 0.001).

### Performance evaluation – Experimental data

The three (N=58 spikes) MU velocity-based signals of human biceps brachii had a firing rate of 11.4 ± 2.6 Hz and an ISI CV of 13.4 ± 0.7% (Table 3). The estimated spike trains from HWM and BPM highly agreed with the EMG reference spike trains (96.3 ± 3.2% and 87.2 ± 10.1%) (Fig. 5A, Table 3). In addition, the spike delta variation for HWM was 1.7 ms, whereas, for BPM, it was 3.7 ms (Fig. 5B).

**Figure 5.**
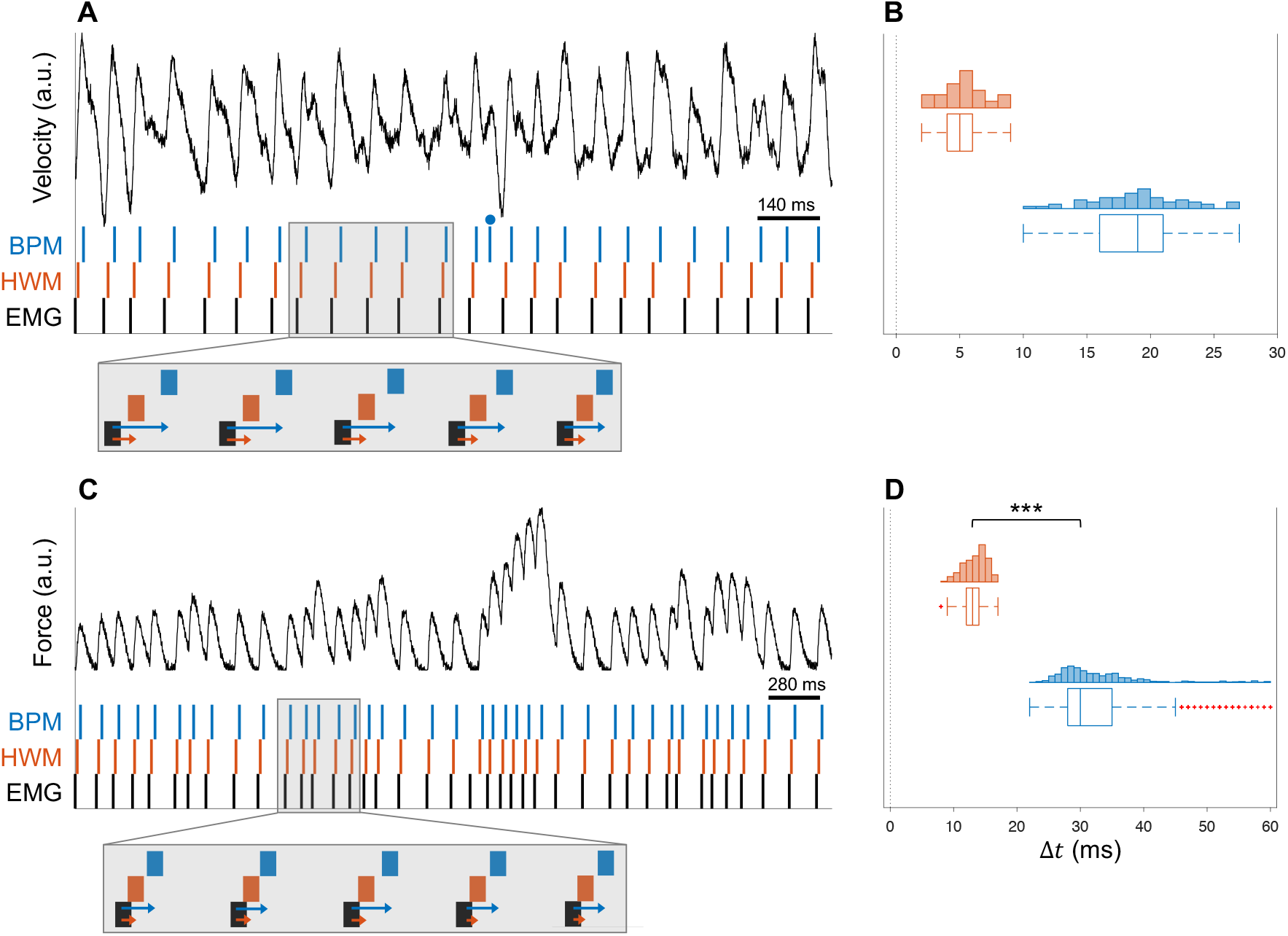
Performance evaluation – Experimental data. **A**. An experimental unfused tetanic signal in terms of velocity (black line) of human biceps brachii during a low force isometric contraction was identified using ultrafast ultrasound. The estimated spikes from HWM (orange vertical lines) and BPM (blue vertical lines) agreed (100% and 96%, respectively) with the concentric needle electromyography reference spikes (black vertical lines). The blue dot denotes a false positive for BPM. **B**. Considering three unfused tetanic contractions of those in **A**, with a total of 58 twitches, the estimated spikes of HWM had a smaller spike delta variation than the estimated spikes by BPM. Note that the mean delay within each experimental dataset was shifted such that the mean was located around 4 ms for HWM and around 20 ms BPM due to subject-dependent electromechanical delay such that comparison of spike estimation variation is possible. **C**. An experimental unfused tetanic signal in terms of force (black line) of rat gastrocnemius during irregular evoked isometric contractions with functionally isolated motor units was measured using a force transducer. The estimated spikes from HWM (orange vertical lines) and BPM (blue vertical lines) agreed (both 100%) with the electromyography reference spikes (black vertical lines). **D**. Considering eighteen unfused tetanic contractions of those in **C**, with a total of 720 twitches, the estimated spikes of HWM had less spike delta and spike delta variation than the estimated spikes by BPM (p < 0.001). Note that the difference in variation is many factors larger than in **B** and that the axis is twice as large.

The eighteen (N=720 spikes) force-based signals of rat gastrocnemius had a firing rate of 14.4 ± 5.5 Hz and an ISI CV of 30.6 ± 2.1% (Table 3). Although the estimated spike trains from HWM and BPM highly agreed with the EMG reference spike trains (96.5 ± 5.6% and 88.0 ± 7.9%) (Fig. 5C), there was a difference in RoA between the two methods (*p* < 0.001) (Table 3). In addition, the average spike delta for HWM and BPM was 12.7 ms and 32.5 ms (Table 3, Fig. 5D). Note the difference in variation is larger than for the velocity-based signals (Fig. 5A-B) due to the different characteristics of the signals. There was a significant difference between the methods’ differences in spike delta (*p* < 0.001) and spike delta variation (*p* < 0.001).

## Discussion

This work aimed to optimize the parameters of HWM and BPM using simulations of unfused tetanic signals. We evaluated the performance of the two methods in terms of the RoA, the spike delta, and the spike delta variability with respect to the simulated/reference spikes. There were three main findings. First, the optimal parameters for HWM are a pseudo-frequency between 45-71 Hz, a minimal peak height between 0.88-0.96, and a minimum peak distance between 15-25 ms. Second, the optimal parameters for BPM are a low-pass cut-off between 20-28 Hz, a minimal peak height between 0.02-0.10, and a minimum peak distance between 10-40 ms. Third, HWM provided estimates with higher RoA, smaller spike delta (bias), and smaller spike delta variation than BPM.

### Simulations

A range of HWM and BPM parameters were robust against various simulation parameters and had a similar agreement (Fig. 3, Fig. S2, Supplementary Material). Although we only considered a limited number of simulation parameters, they should be representative values for low threshold MUs. The upper bound of optimal minimal peak distance values of HWM and BPM was 25 ms and 40 ms, respectively. Those values correspond to a firing rate of 40 Hz and 25 Hz, respectively, which is good enough for lower force levels. The methods will eventually break down by lowering the SNR substantially. Still, we selected the lowest SNR to be 10 dB, which we considered reliable for experimental data, supported by the optimized methods’ high agreement with the EMG references for the 21 experimental signals. Given this, the robustness of the methods’ optimal parameters across these simulation parameters suggests that the parameters not covered in this study would not change the optimal parameters as long as they are within the low threshold MUs setting.

The spike train estimates from HWM had lower spike delta (bias) and spike delta variability than the BPM estimates. The difference in variability is partly explained by the variation in successive twitch shapes, where HWM use the rise of each twitch that should be more robust and vary less across successive twitches than the maximal amplitude that BPM considers (Fig. 2B-C). Another factor we found explaining the spike delta variability was the increase in ISI CV by considering equally shaped twitches (Fig. S3, Supplementary Material). In this study, we also chose the electromechanical delay to be constant and zero. If the value is nonzero but still constant, there will only be a shift in the spike delta (and consequently a modification of the tolerance criteria of the RoA); however, if the electromechanical delay varies within the same contraction. In that case, the spike estimates have an additional variation component, which may be problematic if the absolute exact discharge onset is crucial such as in the analysis of the temporal activity in simultaneous measurements of high-density surface EMG and ultrasound. But varying electromechanical delay within a contraction should not be the case for healthy subjects under normal conditions [23].

We simulated individual twitches using a six-parameter analytical equation [18], where we randomly generated the six parameters from a uniform distribution with pre-defined values (Table 2). These pre-defined values are similar to those estimated in previous work [19]. Different values should not affect this study’s results because the features used by the two spike estimation methods are based on the general twitch shape, i.e., a bell-shaped curve [18], along with a standardised signal. In addition to our simulation model, other models have been suggested in the literature. Raikova et al. proposed a simulation model similar to ours using the same six-parameter analytical equation, but they simplified the model using equally shaped twitches [34]. However, the unfused tetani of voluntary contractions consist of variable successive twitches [16–18]. Fuglevand et al. proposed a model considering recruitment and rate coding in MU pools [20]. Their model considers a twitch model using a power function and two parameters, i.e., the maximal twitch force and the contraction time. Since their model results in a fixed relationship between the maximum twitch force and the contraction time, it is insufficient to describe the considerable variability in real muscle twitches. Moreover, different muscles have MUs with different dynamics of force development and variable twitch parameters that do not fulfil a strict dependence between maximal twitch force and contraction time. In contrast to simulating spike trains and analytical twitch functions, cross-bridge cycling models incorporate Ca^2+^ concentrations [35–38]. For example, Smith et al. used a five-state model of cross-bridge cycling to simulate twitches and unfused tetanic contractions based on a system of differential equations [38]. Although we could have used a more complex model like Smith et al., we are certain that the results should not have changed, and the model we used was adequate for the aims of this study.

### Experimental performance

The estimated spike trains provided by the optimized methods (HWM and BPM) highly agreed with the velocity-based unfused tetanic signals of human biceps brachii and the force-based unfused tetanic signals of rat gastrocnemius during irregular evoked contractions (Fig. 5A and 5C, Table 3). Similar to simulated data, the estimated spikes of HWM had less variation than the estimated spikes by BPM (Fig. 5B and 5D). An interesting finding was that HWM’s estimates provided a spike delta variation that was similar between the velocity-based and the force-based contractions (1.7 ms and 1.8 ms, respectively) but shifted about 8 ms (Table 3). In contrast, for the BPM, the spike delta variation was larger (3.7 ms vs 7.0 ms, respectively) and shifted 14 ms. The larger variation explains the increased spike delta variation in the contraction time parameter (*T*_*c*_), the local maxima that BPM considers in force or displacement signals of twitches. Therefore, the HWM was a better spike estimation method than BPM because it had a higher agreement with reference spikes, lower spike delta (bias), and lower spike delta variability.

### Applications

The applications for a robust spike estimation method would be to get indirect access to the neural drive of the spinal cord via the spinal motor neurons to the muscles by estimating it through the measured unfused tetanic signals. Particularly through the MU-UUS analysis [12], but possibly through the MU-MRI [1,2] or other methods or techniques in the future. Since each spike estimation method (BPM and HWM) took about 2.5 ms to run for an 8-second-long signal consisting of 100 twitches, they are feasible for real-time implementation. In the short term, this method will be important in further developing the identification and analysis of MUs using imaging.

## Conclusions

This study optimized the parameters of a continuous Haar wavelet transform method (HWM) and a band-pass filter method (BPM) using simulations of unfused tetanic signals. We evaluated the performance of the two methods using simulations and two experimental datasets in terms of rate of agreement, spike delta, and spike delta variability with simulated/reference spikes. We found that both methods agreed highly with simulated and reference spikes. However, HWM was a better spike estimation method than BPM because it had a higher agreement, less bias, and less variation. This spike estimation method is important in further developing the identification and analysis of MUs using imaging providing indirect access to the neural drive of the spinal cord via the spinal motor neurons to the muscles by the unfused tetanic signals.

## Acknowledgements

The authors would like to thank Jan Celichowski and Rositsa Raikova for providing excellent quality experimental data on irregularly stimulated single motor units.

