## Supplementary Material for "Optimization and comparison of two methods for spike train estimation in an unfused tetanic contraction of low threshold motor units"

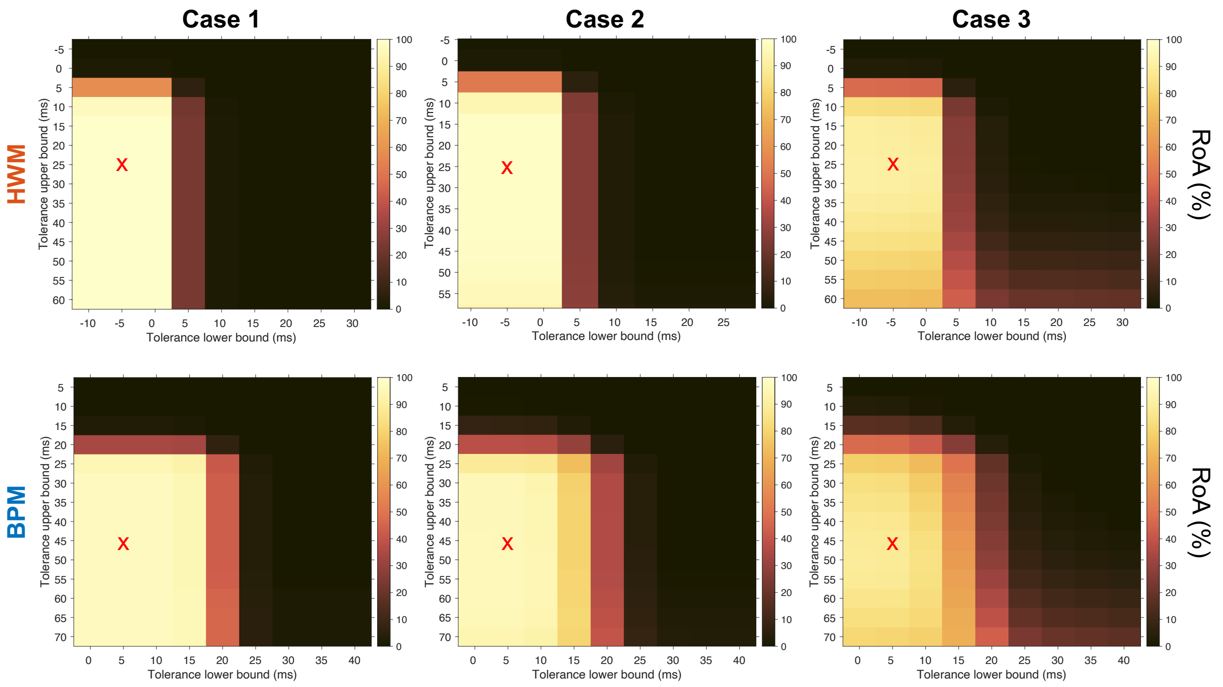


***Figure S1.*** *Sensitivity analysis of the tolerance interval for rate of agreement (RoA) calculation using three cases of 5,000 spikes and subsequent twitches. The first case included a firing rate of 8 Hz, an ISI CV of 5%, and an SNR of 30 dB. The second case included a firing rate of 12 Hz, an ISI CV of 20%, and an SNR of 20 dB. The third case included a firing rate of 16 Hz, an ISI CV of 40%, and an SNR of 10 dB. The red crosses denote the selected tolerance intervals for the optimization. HWM = Haar-wavelet method. BPM = Band-pass filter method.*

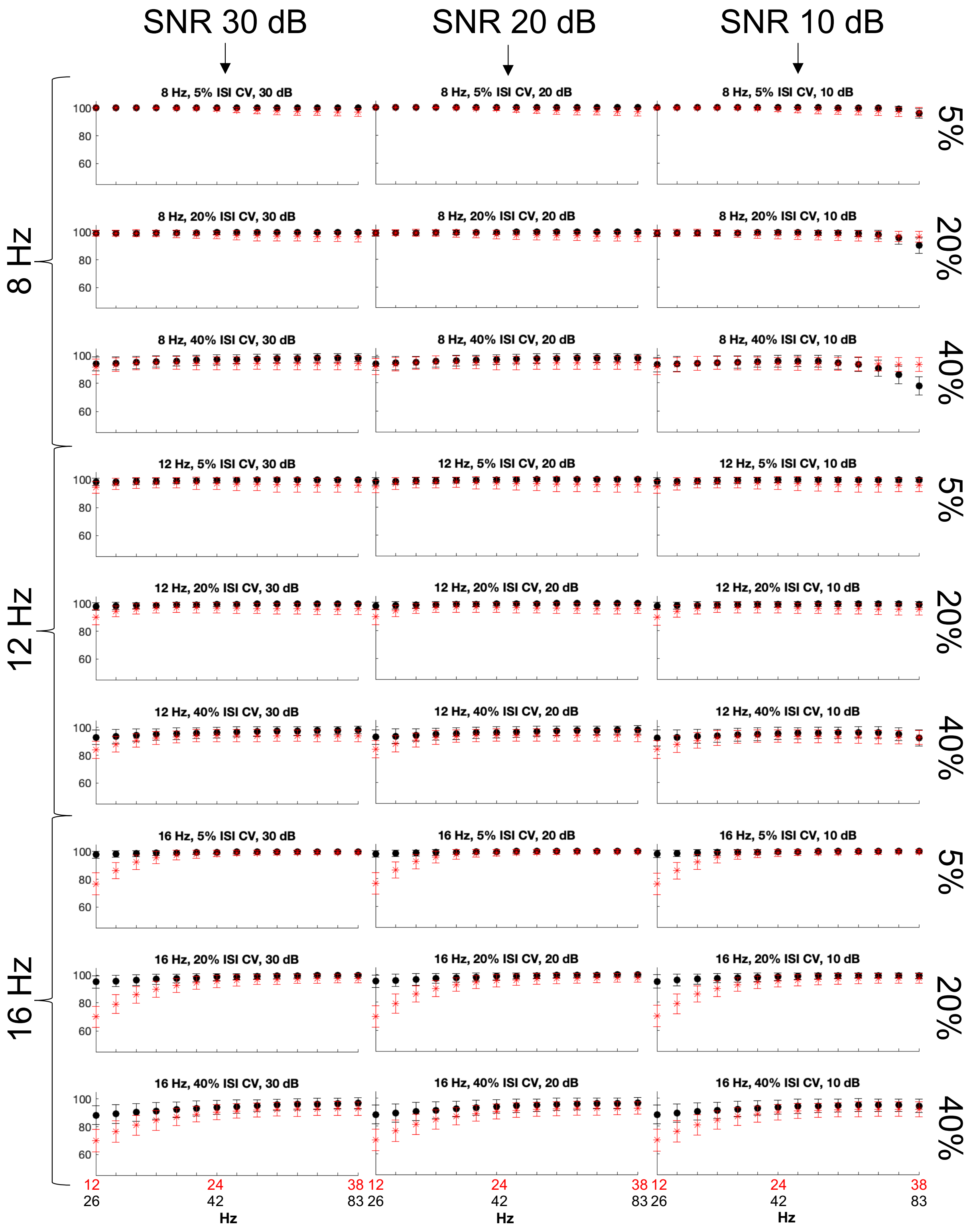


***Figure S2.*** *Quantifying the optimal parameters for each method using simulated data and the rate of agreement (RoA) metric. A total of 270,000 twitches based on different firing rates (8-16 Hz), inter-spike interval coefficient of variations (5-40%), and signal-to-noise ratios (10-30 dB) were considered. The figure shows the mean RoA with respect to different pseudo-frequencies (black dots) concerning the Haar-wavelet method (HWM) and different low-pass cut-off (red asterisks) parameters concerning the band-pass filter method (BPM).*

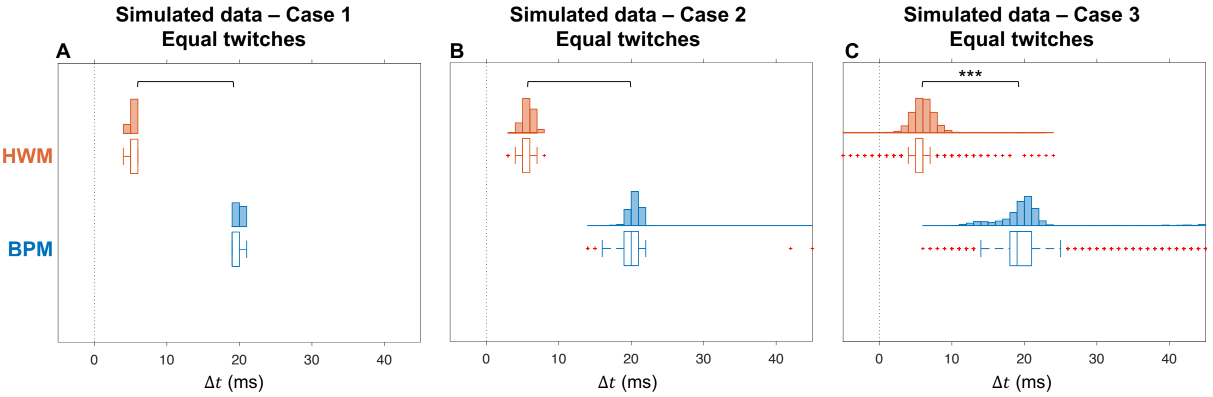


***Figure S3.*** *Performance evaluation – Simulations with equally shaped twitches. Quantifying the method's delay (spike delta) and variation (spike delta variation) between the estimated and the simulated spikes in three cases of complexity (see Fig. S1). The simulated data consisted of 5,000 spikes and subsequent equally shaped twitches.* ***A.*** *Case 1: The estimated spikes of HWM did not have less variation than the estimated spikes by BPM (p = 0.08).* ***B.*** *Case 2: The estimated spikes of HWM did not have less variation than the estimated spikes by BPM (p = 0.15).* ***C.*** *Case 3: The estimated spikes of HWM had less variation than the estimated spikes by BPM (p < 0.001).*
